# Transcriptional signatures of wheat inflorescence development

**DOI:** 10.1101/2022.07.06.498941

**Authors:** Carl VanGessel, James Hamilton, Facundo Tabbita, Jorge Dubcovsky, Stephen Pearce

**Author notes:** Email addresses: Carl VanGessel – James Hamilton – Facundo Tabbita - Jorge Dubcovsky– Stephen Pearce –.

## Abstract

In order to maintain global food security, it will be necessary to increase yields of the cereal crops that provide most of the calories and protein for the world’s population, which includes common wheat (*Triticum aestivum* L.). An important factor contributing to wheat yield is the number of grain-holding spikelets which form on the spike during inflorescence development. Characterizing the gene regulatory networks controlling the timing and rate of inflorescence development will facilitate the selection of natural and induced gene variants that contribute to increased spikelet number and yield.

In the current study, co-expression and gene regulatory networks were assembled from a temporal wheat spike transcriptome dataset, revealing the dynamic expression profiles associated with the progression from vegetative meristem to terminal spikelet formation. Consensus co-expression networks revealed enrichment of several transcription factor families at specific developmental stages including the sequential activation of different classes of MIKC-MADS box genes. This gene regulatory network highlighted interactions among a small number of regulatory hub genes active during terminal spikelet formation. Finally, the *CLAVATA* and *WUSCHEL* gene families were investigated, revealing potential roles for *TaCLE13, TaWOX2*, and *TaWOX7* in wheat meristem development. The hypotheses generated from these datasets and networks further our understanding of wheat inflorescence development.

## INTRODUCTION

The world population is expected to exceed nine billion people by 2050, signaling that further increases in grain production will be required to ensure food security ^1^. Because there remain few opportunities to expand arable land area, increasing the yield of major cereal crops through genetic improvement will be critical to meet this goal. In common wheat (*Triticum aestivum* L.) characterizing the genetic pathways regulating grain size and grain number will facilitate the rational combination of superior alleles in wheat breeding programs to help drive continued yield improvements ^2^.

Grain number in wheat is determined to a large extent by inflorescence architecture. By integrating photoperiod and temperature cues, the vegetative shoot apical meristem (SAM) transitions to the reproductive inflorescence meristem (IM), during which the developing spike passes through the characteristic double ridge (DR) stage, forming a lower leaf ridge and an upper spikelet ridge ^3^. The lower leaf ridge is repressed by the MIKC-MADS box transcription factors (TFs) *VRN1, FUL2* and *FUL3* ^4^, whereas the upper ridges develop glumes, lemmas, and floret primordia. As the IM elongates, spikelet meristems are added at the growing apex, while basal spikelets continue to develop. Wheat spikes are determinate structures and the addition of lateral spikelets ends when the terminal spikelet is formed. Therefore, spikelet number is determined by the timing and rate of meristem development preceding terminal spikelet formation. Each spikelet has the potential to form between three and six grains ^5^ and spikelet number is correlated with grain number and yield ^6–8^.

Shoot meristems are organized around the organizing center and stem cell maintenance is governed by the conserved CLAVATA-WUSCHEL negative feedback loop ^9^. In Arabidopsis, the homeodomain TF *WUS* induces *CLV3*, which encodes a secreted peptide that forms receptor complexes repressing *WUS* ^10^. Manipulation of this pathway confers variation in locule number in tomato (*Solanum lycopersicum*) and kernel row number in maize (*Zea mays*) ^11,12^. The wheat genome contains 104 *CLAVATA3/EMBRYO SURROUNDING REGION* (*CLE*) peptides ^13^ and 44 WUSCHEL RELATED HOMEOBOX (WOX) TFs^14^, but the specific ones regulating inflorescence meristem development in wheat are yet to be identified.

Inflorescence development is controlled by a complex regulatory network involving multiple classes of transcription factors (TFs) which orchestrate rapid and dynamic changes in gene expression. The Type II MIKC MADS-box TFs play critical roles in flower development across the angiosperms and can be divided into A, B, C, D and E-classes that interact mainly as tetrameric complexes in a spatially regulated manner to direct sepal (A- and E-), petal (A-, B-, E-), stamen (B-, C-, E-), and carpel development (C- and E-class genes) ^15,16^. This family expanded during cereal evolution and the hexaploid wheat genome contains 201 MIKC MADS-box genes, classified into 15 phylogenetic subclades ^17^.

The SHORT VEGETATIVE PHASE (SVP) subclade members *SVP1, VRT2*, and *SVP3* promote the transition from the vegetative SAM to the IM, along with the AP1/SQUA subclade genes *VRN1, FUL2* and *FUL3* ^4,18^. Subsequently, AP1/SQUA genes suppress the expression of SVP genes, which may be required to promote interactions between AP1/SQUA proteins and the E-class MIKC-MADS proteins SEPELLATA1 (SEP1) and SEP3, which are predominantly expressed in floral organogenesis during early reproductive growth ^18^. The natural *VRT2*^*pol*^ allele from *Triticum polonicum* exhibits ectopic expression and is associated with elongated glumes and increased grain length ^19^. *VRT2*-overexpression lines show reduced transcript levels of B-class (*PI* and *AP3*) and C-class (*AG1* and *AG2*) MIKC-MADS box genes, although the role of these latter subclades in wheat inflorescence development remains to be characterized ^18^.

Although much has been learned about wheat inflorescence development from positional cloning, reverse genetics and comparative genetic approaches, we lack a full understanding of the regulatory networks controlling meristem determinacy and developmental transitions. Only a fraction of the hundreds of QTL for thousand kernel weight, kernel number per spike, and spikelet number have been cloned and validated to date, indicating that a large proportion of quantitative variation in these traits remains uncharacterized^7^.

Transcriptomics provides a complementary approach to characterize the regulatory networks underlying inflorescence development that is empowered by an expanding set of wheat genomic resources^20,21^. Co-expression and gene regulatory networks (GRNs) are powerful tools to interpret temporal correlation and causal relationships between genes, and to help identify critical hub genes that coordinate development ^22,23^. Previous transcriptomic studies in wheat inflorescence tissues described the differential expression profiles of thousands of genes during vegetative and floral meristem development, including the stage-specific expression of different TFs and hormone biosynthesis and signaling genes ^24,25^. A population-associative transcriptomic approach was used to identify regulators of wheat spike architecture, including *CEN2, TaPAP2*/*SEP1*-*6*, and *TaVRS1*/*HOX1*, which were validated in functional studies ^26^.

In the current study, a series of co-expression and gene regulatory networks were assembled to characterize the predominant transcriptional profiles associated with the progression of wheat inflorescence development, revealing two consecutive regulatory shifts at the DR and TS stages. Core regulatory candidate genes were identified including both known TFs and novel candidates with potential roles in regulating spike architecture.

## RESULTS

### Early wheat inflorescence development is defined by two major transcriptional shifts

To characterize the wheat transcriptome during inflorescence development, RNA was sequenced from tetraploid durum wheat meristem tissue at five developmental stages; vegetative meristem (W1), double ridge (W2), glume primordium (W3.0), lemma primordium (W3.25), and terminal spikelet (W3.5) (Fig. 1A) ^3^. An average of 28.9 M reads per sample (79.6% of all reads) mapped uniquely to the A, B and U genomes of the IWGSC RefSeqv1.0 assembly. Of the 190,391 gene models on these chromosomes, 82,019 (43.1%) were expressed (> zero TPM) and 45,243 (23.8%) had a mean expression greater than one TPM in at least one timepoint (Supplementary data 1). Of the 3,861 gene models annotated as TFs (2.0% of gene models), 2,874 (74.5%) were expressed (> zero TPM) and 1,703 (44.1%) had a mean expression greater than one TPM in at least one timepoint (Supplementary data 2).

**Figure 1:**
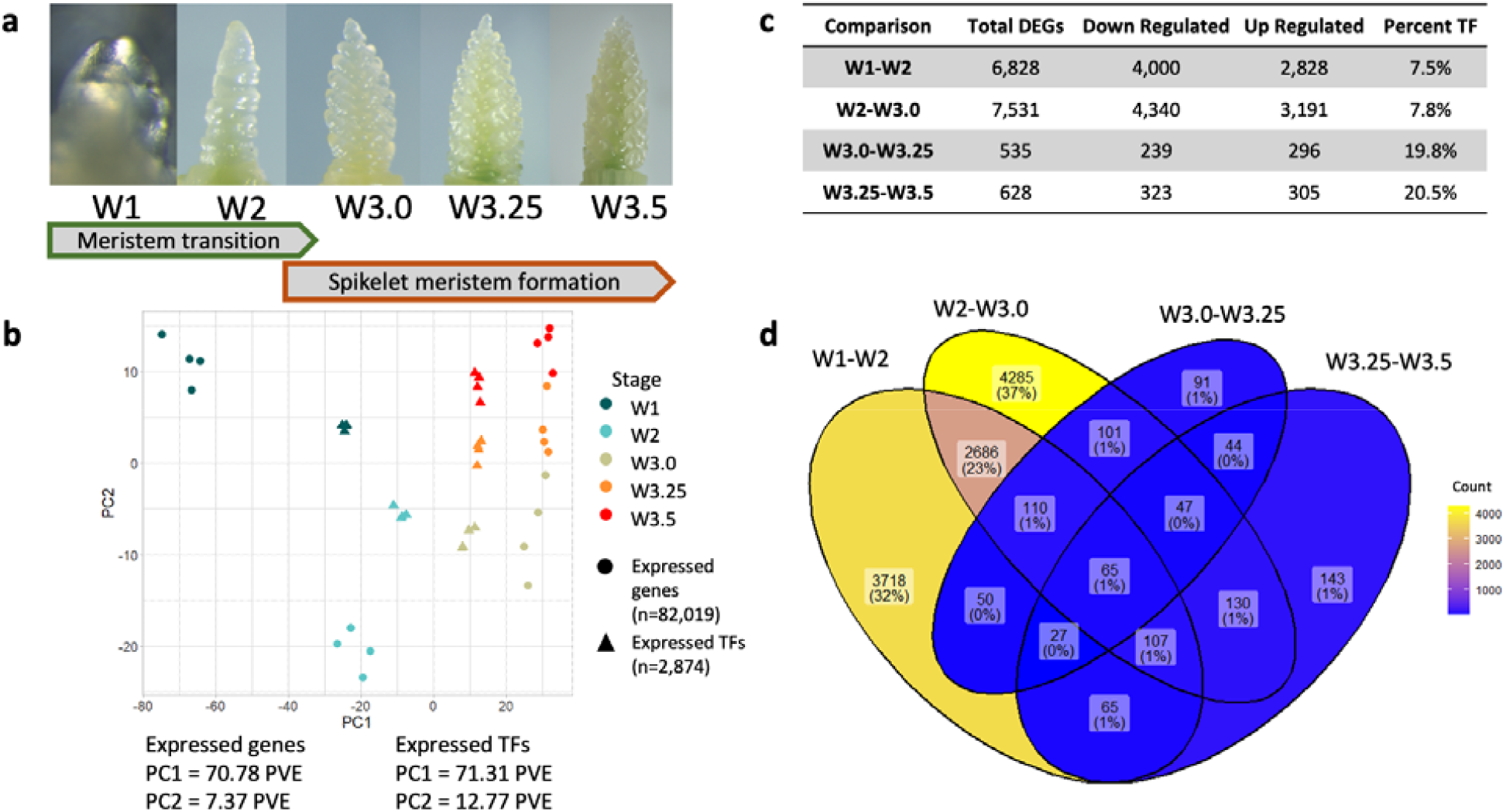
The early wheat inflorescence development transcriptome. (A) Sampling stages of Kronos apical meristems according to the Waddington development scale^3^; W1.0 – vegetative meristem, W2 – double ridge, W3.0 – glume primordium, W3.25 – lemma primordium, W3.5 – terminal spikelet. (B) Whole transcriptome and transcription factor expression principal component analysis of samples, PC1 plotted on the x-axis and PC2 plotted on the y-axis. PVE = Percent Variance Explained. (C) Differentially expressed genes (DEGs) in sequential pairwise comparisons (W1 – W2, W2 – W3.0, W3.0 – W3.25, W3.25 – W3.5). The total number of genes, the number up- and down-regulated and the proportion encoding transcription factors (TF) are described. (D) Venn diagram of DEGs in each consecutive pairwise comparison from (C). Each category is shaded according to the number of sequential DEGs shared among the four comparisons.

Comparison of the inflorescence development transcriptome with two whole-plant wheat development transcriptome datasets ^27,28^ revealed 3,682 genes with spike-dominant expression profiles (τ > 0.9, where zero means constitutive expression and one indicates tissue-specific expression) (Supplementary data 3). These genes were most strongly enriched for gene ontology (GO) terms relating to histone assembly and chromosome organization (Supplementary data 4), but also included 286 genes (7.8%) encoding TFs, including both *LEAFY* homoeologs, 15 GROWTH REGULATING FACTOR (GRF) TFs (of 20 expressed during the time course), seven SHI RELATED SEQUENCE (SRS) TFs (out of ten), 20 TCP TFs (out of 49) and ten WOX TFs (out of 28, Supplementary data 3). Despite their known roles in regulating inflorescence development, only two out of 130 MIKC-MADS box and six out of 41 SPL TFs exhibited spike-dominant expression profiles, suggesting they play more diverse roles across plant development. There were 86 spike-specific genes with zero expression in all other stages of development (τ = 1) (Supplementary data 3).

Principal component analysis (PCA) using the whole transcriptome grouped the four biological replicates of each growth stage closely together and revealed that the majority of the transcriptional changes in this time course occur between the vegetative meristem and double ridge formation (Fig. 1B). These changes are described by PC1, which accounted for 71.8 percent variation explained (PVE). The transition from W1 to W2 was associated with 6,828 DEGs, 58.6% of which were downregulated (Fig. 1C, Supplementary data 5) and most significantly enriched for GO terms relating to “cell wall organization”, and lignin and hemicellulose metabolic processes (Supplementary data 6). Surprisingly, the 2,828 (41.4%) DEGs upregulated between W1 and W2 were most significantly enriched for GO terms relating to photosynthesis despite the transition from leaf to floral meristem development (Supplementary data 5).

The transition from W2 to W3.0 was associated with 7,531 DEGs (57.6% downregulated, Supplementary data 5, 6). The 3,191 DEGs upregulated between these timepoints were most significantly enriched for “meristem maintenance” and “flower development” GO terms (Supplementary data 6), suggesting that a number of genes triggering floral meristem formation are first activated at this stage.

By contrast, the transcriptomic changes from W3.0 to terminal spikelet formation (Fig. 1A) were distributed across PC2, which accounts for just 7.4 PVE (Fig. 1B) and were associated with 12.3-fold fewer DEGs than during the transition from vegetative meristem to stage W3.0 (Fig. 1C). Just 535 DEGs were found between W3.0 and W3.25 (55.3% upregulated) and 628 DEGs between W3.25 and W3.5 (48.6% upregulated) (Supplementary data 5). Genes upregulated across these three timepoints were most significantly enriched for “floral organ identity” (Supplementary data 6). There are fewer developmental changes between W3.25 and W3.5, relative to changes between W1 and W3.0, which may be due in part to basal and apical spikelets being at similar developmental stages between the latter timepoints ^29^.

Of the 11,669 DEGs in at least one of the four consecutive pairwise comparisons, 899 (7.7%) encoded a TF, a 2.2-fold enrichment (hypergeometric *P* = 2.22 e-62). This enrichment was strongest after DR through terminal spikelet formation (5.2-fold enrichment, *P* = 8.73 e-73) where TFs accounted for 19.8% and 20.5% of all DEGs in pairwise comparisons (Fig. 1C). A PCA using only TF expression resulted in the same spatial arrangement of biological samples as in the whole-transcriptome PCA but with improved resolution between stages (Fig. 1B), and explained a greater proportion of variation for PC2 than when including the whole transcriptome (Fig. S1).

Taken together, these analyses show that less than half of the wheat transcriptome but nearly three-quarters of TFs are expressed during inflorescence development, including a set of genes which are spatially and temporally restricted to early inflorescence tissues. Terminal spikelet formation is associated with comparatively less transcriptional variation relative to stages preceding W3.25 and the strong enrichment in TFs suggests they play critical roles during this stage.

### Co-expression networks reveal predominant transcriptome profiles during inflorescence development

Co-expression networks were assembled to identify highly correlated modules of genes that define the major transcriptional profiles during early inflorescence development. All networks were assembled using a set of 22,566 genes that were differentially expressed in at least one of the ten possible pairwise combinations between timepoints (Fig. 1D) and that were also defined as significantly differentially expressed using ImpulseDE2, a package used to analyze longitudinal transcriptomic datasets (Supplementary data 5).

A consensus network constructed with repeated subsampling and randomized parameters with WGCNA (see Materials and Methods) assembled these genes into 21 modules with a mean connectivity score of 0.485 (Fig. 2A, Supplementary data 7). A standard WGCNA network was also constructed using ‘best practices’ parameters but with no repeated subsampling and randomization and had a connectivity score of 0.327 which skewed to zero (Fig. 2A). In both networks, the majority of genes clustered into modules 1 and 2, which contained many of the same genes (Jaccard index > 0.86, Fig. S2). However, other modules exhibited dissimilar expression profiles between networks (Jaccard index < 0.5), indicating the consensus network clustered genes into a greater number of modules with distinct expression profiles not captured in the standard network. Based on the improved correlation of co-clustered genes within modules and the detection of distinct regulatory profiles, the consensus network was used in all subsequent analyses.

**Figure 2:**
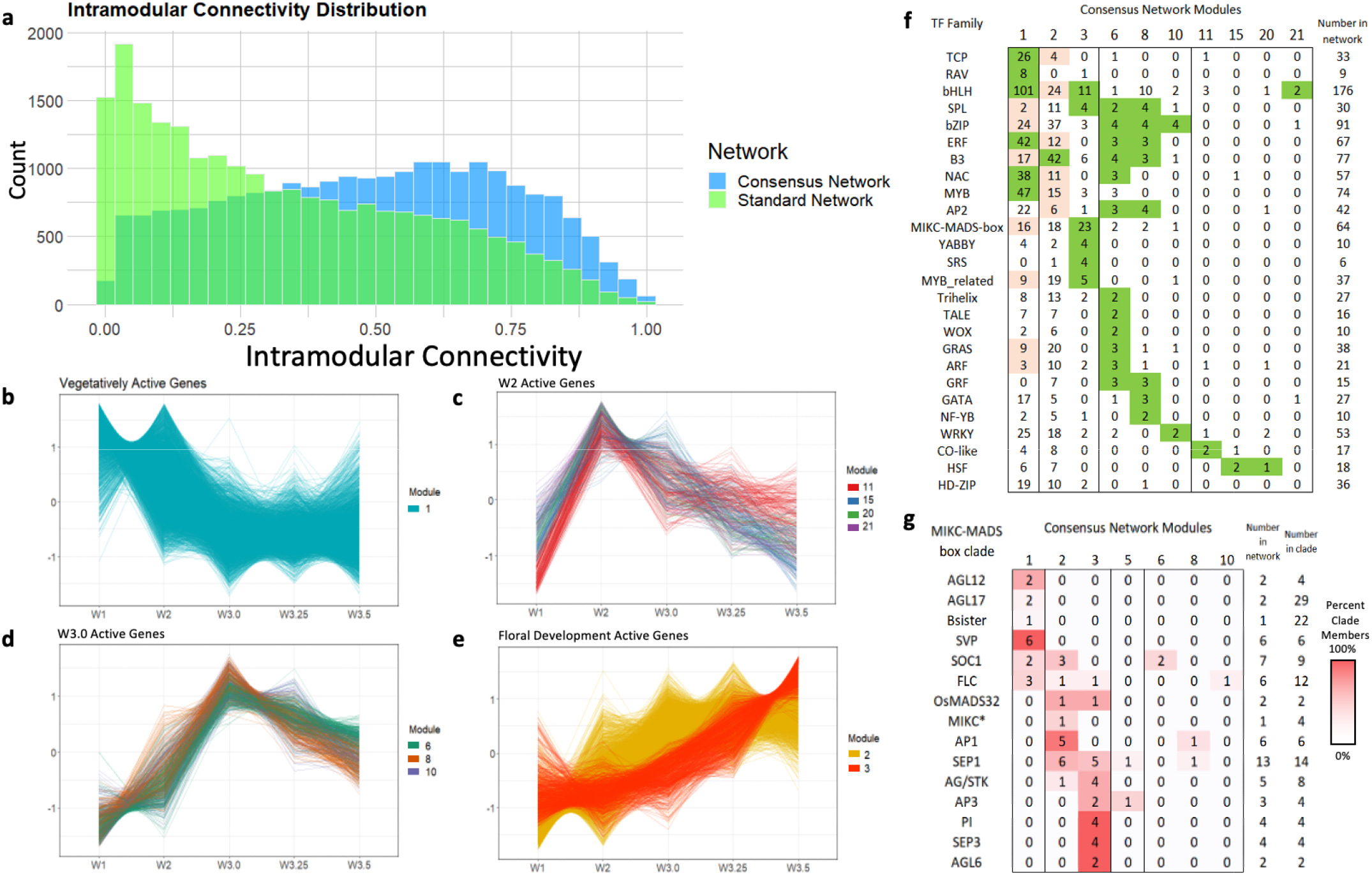
Co-expression networks showing the dominant transcriptional profiles during wheat inflorescence development. (A) Histogram of intramodular connectivity scores for 22,566 genes clustered in consensus (blue) or standard (green) network. (B - E) Expression profiles during inflorescence development of discussed modules in the consensus network. Lines represent scaled time course expression of each gene in the module. Modules with similar expression profiles are grouped together for comparison. (F) Number of TF family members clustered in each discussed consensus module. Modules enriched (green) or depleted (pink) for TF families are highlighted (*P* < 0.01). (G) Number of MIKC-MADS box clade members clustered in each consensus modules. Co-expressed MIKC-MADS box groups are shaded relative to the total number of genes in the clade.

### Inflorescence meristem development is associated with the down-regulation of RAV and TCP transcription factors

Module 1 was the largest in the network and grouped 10,102 genes defined by high transcript levels in the vegetative meristem and early meristem transition followed by down-regulation after DR and as the spike develops (Fig. 2B). Several TF families were enriched in this module, including 101 basic Helix-Loop-Helix (bHLH) TFs, 47 MYB TFs and eight of the nine differentially expressed RELATED TO ABI3 AND VP1 (RAV) TFs included in the network (Fig. 2F). Twenty-six of the 33 total TCP TFs clustered in this module, nine of which were also spike-dominant expressed (Fig. 2F). Although at the whole family level MIKC-MADS TFs are significantly under-represented in module 1 (Fig. 2F, hypergeometric *P* = 8.6 e-4), all six SVP genes (*SVP1, VRT2* and *SVP3*) cluster in this module, consistent with their specific role regulating early stages of inflorescence development. In addition, both AGL12 subclade genes, and three of the six FLC subclade genes clustered in this module (Fig. 2G).

### A small number of genes are transiently expressed during double ridge formation

Genes which showed a peak at the double ridge stage (W2) followed by a decline in later stages were clustered in modules 11 (131 genes), 15 (104 genes), 20 (44 genes) and 21 (42 genes). These clusters share broadly similar expression profiles (Fig. 2C) and were enriched for genes with spike-dominant expression profiles (between 2.1 and 3.0-fold enrichment). Genes in modules 15 and 20 were significantly enriched for development functional terms including “shoot system development” and “carpel development” (Supplementary data 8) including three *TERMINAL FLOWER1-like* genes *CENTRORADIALIS2* (*CEN2*), *CEN4*, and *CEN-5A* (Supplementary data 7). All three modules were enriched for the functional term “response to auxin” and included several auxin-responsive factors (ARF), indole-3 acetic acid (IAA), and SAUR-like protein family members, indicating that auxin signaling may promote double ridge formation.

### Inflorescence transition and spike architecture genes are upregulated at W3.0

Modules 6 (267 genes), 8 (211 genes), and 10 (144 genes) share broadly similar profiles defined by maximum expression at stage W3.0 and subsequent downregulation (Fig. 2D). Each of these modules was significantly enriched (between 2.3 and 5.3-fold) for spike-dominant genes, indicating they likely play highly specific roles restricted to developing meristems and inflorescence initiation. Module 6 included 18 genes previously associated with variation in spikelet number and five orthologs of rice genes with roles in panicle development, including the ERF TF *WHEAT FRIZZY PANICLE* (*WFZP*) and *KAN2*, a MYB TF which functions in establishing lateral organ polarity in *Arabidopsis* ^30,31^.

### Inflorescence and spikelet meristem formation is associated with sequential activation of different classes of TFs

The 8,971 genes in module 2 were defined by the inverse transcriptional profile to module 1, with low expression in the vegetative meristem followed by sustained upregulation from the double ridge stage onwards (Fig. 2E). Transcription factors were under-represented in this module, and only the B3 family (42 of 77 B3 TFs assembled in the co-expression network) was significantly enriched (Fig. 2F). There were 18 MIKC-MADS box TFs which were upregulated early in the transition to the inflorescence meristem including all genes in the AP1/SQUA subclade (with the exception of *VRN-A1*) and six of the thirteen genes in the SEP1 subclade (Fig. 2G). Several genes with characterized roles in inflorescence development clustered in this module, including *FLOWERING LOCUS T2* (*FT-A2*), *Q*, and *RAMOSA2* (*TaRA-B2*) (Supplementary data 7)^32,33^.

The 708 genes clustered in module 3 exhibited a similar transcriptional profile to module 2, with a delayed upregulation and stronger peak at the terminal spikelet stage (Fig. 2E). These genes are significantly enriched for developmental functional terms including “specification of floral organ identity”, suggesting they include floral patterning and developmental genes that regulate spikelet meristem formation (Supplementary data 8). This module was significantly enriched for both spike-dominant expressed genes (106 genes, *P* < 0.001) and for TFs (86 genes, 12.1%, *P* < 0.001), consistent with pairwise DE analysis between stages W3.0 and W3.5 (Fig. 1C). These included four members of the SRS TF family, four YABBY TFs, and the HD-zip TFs *Grain Number Increase 1* (*GNI1*) and *HOX2* (Supplementary data 7). All members of the MIKC-MADS subclades PI, AGL6 and SEP3 were clustered in module 3, as well as two of the three AP3 subclade genes, four of the five AG/STK subclade genes and five SEP1 subclade genes (Fig. 2G).

### Gene regulatory networks predict high-confidence interactions between transcription factors

To identify the most robust co-expression patterns, the consensus adjacency matrix used for previous co-expression analyses was filtered for genes which co-clustered with at least one gene every time they were co-sampled in 1,000 networks assembled with variable, randomized parameters. The 18,174 genes that met this criterion were assembled into a conensus100 network consisting of 924 modules with a median size of three (Supplementary data 7).

Module 9 of this network comprised 167 genes (including 32 TFs) which were most highly expressed at the terminal spike stage (Fig. S3) and significantly enriched for the GO terms “specification of floral organ identity” and “flower development” (Supplementary data 9), suggesting it may represent a core regulatory network for wheat spikelet and/or floret development. The genes with the highest connectivity (Kw, a measure of each gene’s intramodular co-expression) in this module are *SEP1-A2* and *SEP1-B2*, which may be related with the intermediate position of the *SEP* genes between the meristem identity SQUAMOSA MADS-BOX genes and the anther and carpel development MADS-box genes. This module also groups *WAPO-A1*, that influences spikelet number and stamen identity ^34^ and a gene encoding an F-box protein that is a component of an SCF ubiquitin ligase that may be targeted by *TB1* ^35^ (Supplementary data 7).

To predict interactions between TFs during inflorescence development, a *de novo* Causal Structure Inference (CSI) network was constructed using all 970 TFs from the consensus100 network. This gene regulatory network consisted of 704 genes (nodes) with 5,604 predicted interactions (edges) with interaction strength (edge weight) > 0.001 (Supplementary data 10). To prioritize the most important regulatory candidate genes, the network was screened for interactions with an edge weight ≥ 0.03, leaving 88 genes with 177 interactions. The majority of these genes were from consensus modules 1 (37 genes, 42.0%) and 3 (36 genes, 40.9%), with 27 of the latter genes clustered in consensus100 module 9 (Fig. 3).

**Figure 3:**
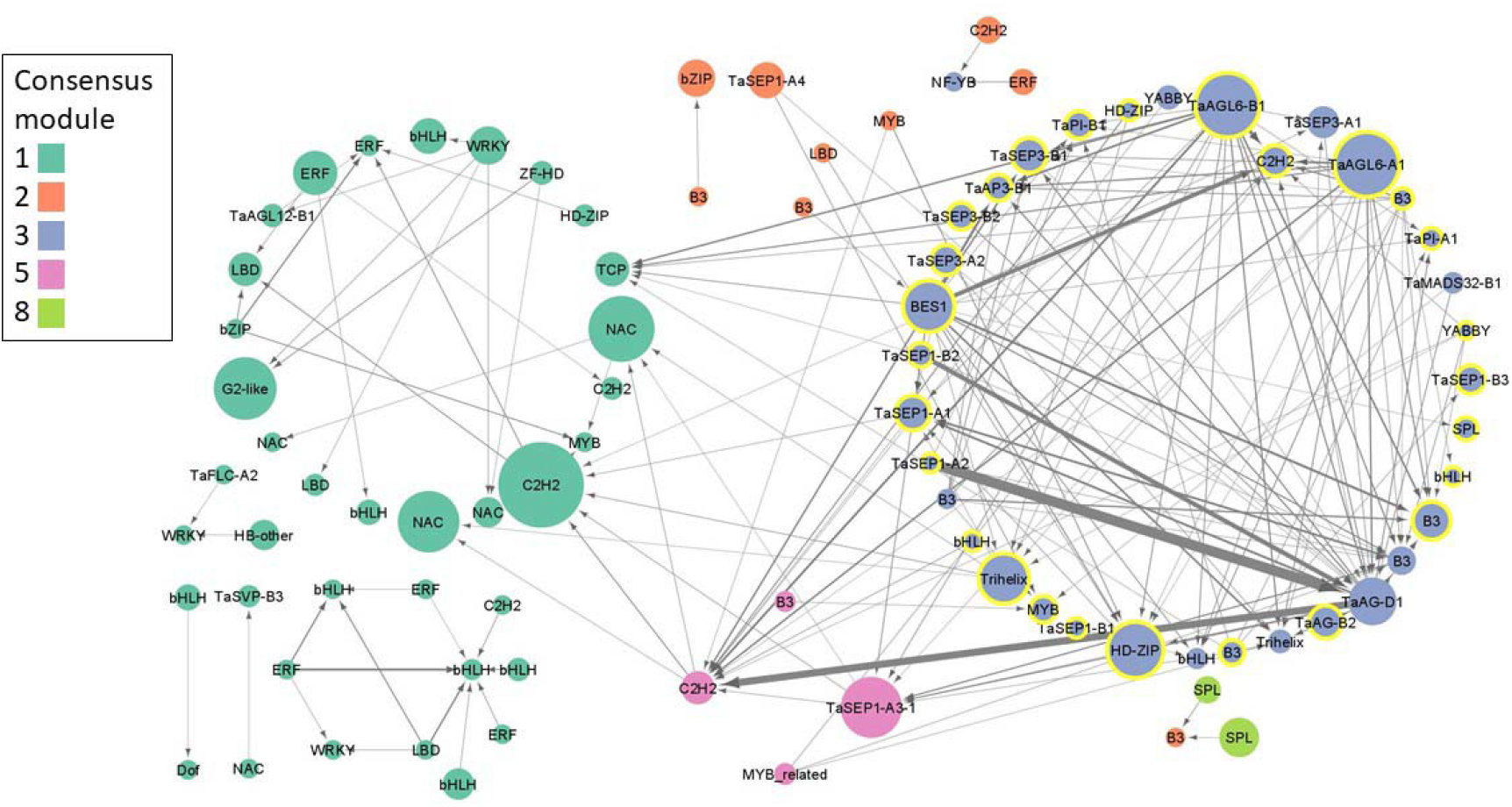
Causal structural inference prediction of interacting transcription factors, filtered for edge weight ≥0.03. Nodes (genes) are colored by their consensus network modules, and consensus100 module 9 genes are highlighted with a yellow border. Node diameter is scaled to betweenness centrality to indicate its importance within the network. Directional interactions are indicated by arrows and width is scaled to predicted interaction strength.

Most predicted interactions were between genes in the same consensus module, with the majority occurring within module 3 and involving MIKC-MADS box TFs, suggesting a closely coordinated network during spikelet meristem and terminal spikelet formation (Fig. 3). Among the genes with the highest betweenness centrality, a measure of each gene’s importance in the overall network, were *AGL6-A1* and *AGL6-B1* which were predicted to interact with 31 other TFs in the network, including 13 MIKC-MADS genes such as *PI-1, SEP3-1, AP3-1, SEP1-1* and *AG1* (Fig. 3). Interaction strengths implicated a role for *AG-D1* as a regulatory hub with strong incoming interactions from other MIKC-MADS-box genes from the SEP1, SEP3, AG, PI, and AP3 subclades, as well as outgoing interactions with genes such as the LOFSEP MIKC-MADS box TF *SEP1-1* (Fig. 3). The BES1 TF *BES1/BZR1 HOMOLOG 2-like* had high betweenness centrality and was predicted to have outgoing interactions with MIKC-MADS, Trihelix and HD-ZIP TFs (Fig. 3).

Cross-module interactions included 16 outgoing edges from module 3 to module 1, including six outgoing interactions to a PCF-type TCP TF (Fig. 3). Although only four TFs from module 5 were assembled in the network, they included *SEP1-A3* and a C2H2 TF with ten incoming interactions from module 3 including *AGL6-B1, BES1/BZR1 HOMOLOG 2-like* and *AG-D1* (Fig. 3).

### Integrating transcriptomics to prioritize candidate genes underlying natural variation

The consensus network includes 4,637 high confidence homoeologous gene pairs, the majority of which (3,636, 78.4 %) clustered either in the same module, or in modules with highly similar expression profiles (Supplementary data 7). We hypothesized that homoeologous genes clustering in different modules may have divergent expression profiles resulting from natural variation in one homoeolog. Of these 1,001 divergently expressed gene pairs, 221 encoded TFs, including *VRN1* (where the dominant *VRN-A1* spring allele is expressed at an earlier stage of inflorescence development compared to the wild-type *VRN-B1* allele), *RHT1* (where the *Rht-B1b* semi-dwarfing allele is more highly expressed in the vegetative meristem than *RHT-A1*), and *TEOSINTE BRANCHED 1* (*TB1*, where *TB-B1* expression is maintained at higher levels than *TB-A1* during terminal spikelet formation, Figure 4A).

**Figure 4:**
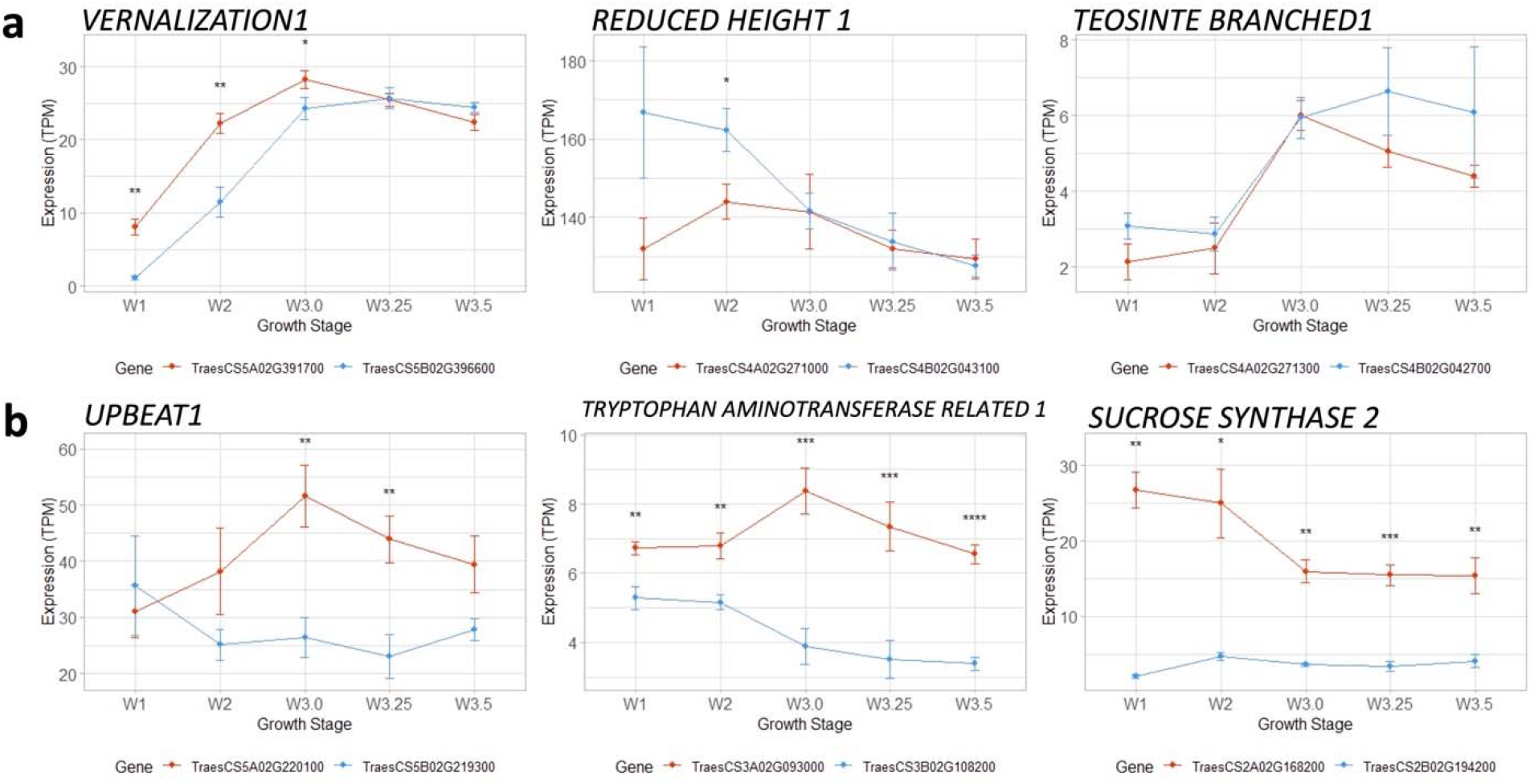
Divergent expression of homoeologous gene pairs during inflorescence development. Expression profiles of (A) Characterized domestication and adaptation alleles and (B) Genes close to QTL for spike architecture or grain size. Expression values are in TPM ± standard error. A-genome homoeologs are in orange, B-genome homoeologs are in blue. Paired t-tests were used to indicate differences between homoeolog expression at each time point, *P* values < 0.05 (*), 0.01 (**), 0.001 (***), 0.0001 (****).

Each of the three genes from Fig. 4A lies within 250 kb of a QTL for either grain number or grain size (Supplementary data 7), so we hypothesized that other differentially expressed homoeologs located close to a yield-component QTL might point to natural variation for yield traits in wheat. For example, *UPBEAT-A1* is upregulated at the double ridge stage to a much greater degree than *UPBEAT-B1* (Fig. 4B), is close to a QTL for TKW, and encodes an ortholog of a bHLH TF that regulates cell proliferation in Arabidopsis ^36^. Similarly, *TRYPTOPHAN AMINOTRANSFERASE RELATED-A1* (*TAR-A1*) is also upregulated at the double ridge stage compared to *TAR-B1* (Fig. 4B) and is proximal to a QTL for grain yield (Supplementary data 7). These genes encode enzymes in the IAA biosynthesis pathway and their overexpression has previously been shown to modify inflorescence development in wheat ^37^. Co-expression networks and observations from meta-analysis are available for developing hypotheses on inflorescence development (Supplementary data 7).

### Identification of CLE/WOX genes expressed during wheat inflorescence development

To identify members of the conserved CLAVATA-WUSCHEL pathway that may regulate stem cell maintenance in wheat spike meristems, the expression profiles of genes encoding WOX TFs and *CLE* peptides were analyzed. Of 29 WOX TFs, 28 were expressed during early inflorescence development and 11 were both significantly differentially expressed during the time course and exhibited a spike-specific expression profile (Fig. 5A). Two orthologs of *OsWOX4* were co-expressed in module 1 with rapid down-regulation before transition to the inflorescence meristem, suggesting they may play a role in vegetative meristem maintenance but not in inflorescence development. Seven *WOX* genes clustered in module 2, characterized by rising expression during inflorescence development, including the orthologs of *AtWUS* (*TaWUSa* and *b*). The homoeologues *TaWOX2a* and *2b* are both associated with variation in spikelet number and are clustered into separate co-expression modules (Supplementary data 7). Of the 64 *CLE* genes, 35 were expressed during inflorescence development and just nine were differentially expressed across the time course (Fig. 5B). Three wheat genes orthologous to *OsFON2/4* (putatively *TaCLV3, TraesCS2A02G329300* and *TraesCS2B02G353000*) exhibit spike-dominant DR-peaking expression profiles.

**Figure 5:**
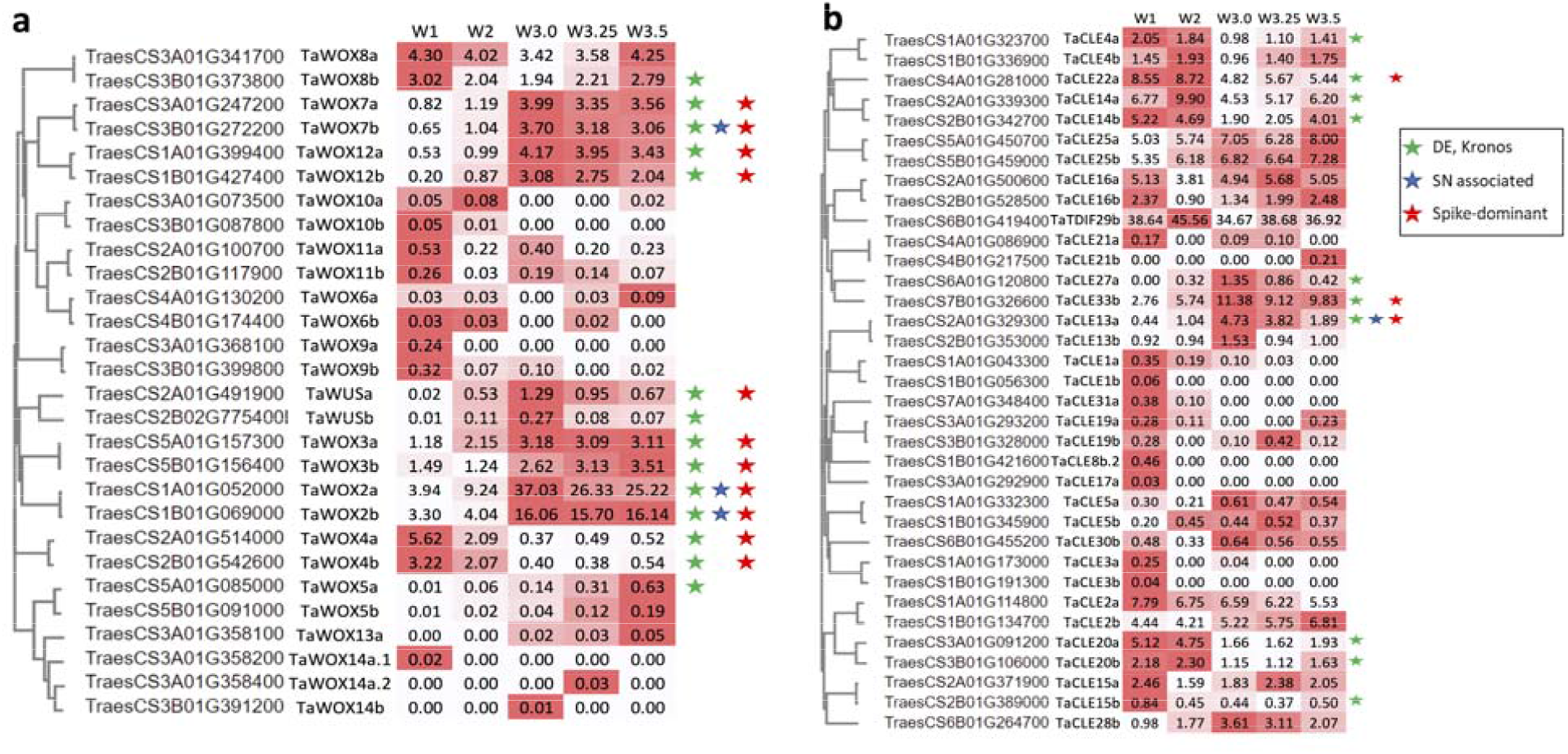
Expression profiles of WOX TFs (A) and CLE peptides (B) during wheat inflorescence development. Stars indicate additional evidence of a possible role in spike regulation (green = differential expression in ‘Kronos’ inflorescence, blue = associated with variation in spikelet number, red = spike-dominant expression profile). Heatmaps show expression (TPM) relative to each gene’s minimum and maximum expression. Only genes with TPM ≥ 0.05 are shown.

## DISCUSSION

Temporal transcriptomic datasets can help to characterize the regulatory networks controlling the development of complex organs such as the wheat inflorescence. One strategy to reduce spurious co-clustering of genes is to assemble a consensus co-expression network using a matrix of co-clustering frequencies from multiple independent networks, each assembled with randomized parameters and gene selection ^38–40^. Co-expression networks have been successfully applied to unravel gene function in yeast (*Saccharomyces cerevisiae*), floral and fruit developmental pathways in strawberry (*Fragaria vesca*), and regulatory networks underlying leaf development in maize (*Zea mays*) ^39–41^. In the current study, this approach generated a consensus network with a larger number of modules with improved intramodular connectivity compared to a standard WGCNA network (Fig. 2A). A further refinement to screen for genes co-clustering in every network assembly that they were both included revealed a consensus100 module 9 of 167 genes that likely contribute to spikelet meristem and terminal spikelet formation (Fig. S3), indicating that consensus networks can help improve the accuracy of co-expression predictions and module assignment.

Beyond co-expression profiles, context-specific gene regulatory networks provide information on the centrality of each gene (a measure of its importance to the flow of information through a network), as well as the strength and directionality of interactions between individual genes ^42^. This network predicts that the MIKC-MADS box TF *AGL6* is a critical gene in inflorescence development regulatory networks, and functions together with MIKC-MADS TFs from the PI and SEP subclades (Fig. 3). This is consistent with its role in rice, where AGL6 functions as a cofactor with A, B, C, and D class proteins during floral development, as well as in wheat, where it interacts with ABCDE proteins, likely as a bridge in complex protein-protein interactions to regulate whorl development ^43–45^. This network also revealed novel candidate genes for future characterization studies. For example, the *BES1* TF *BES1/BZR1 HOMOLOG 2-like* is predicted to interact with several TFs, including two HD-ZIP TFs with homology to *HvVRS1*, suggesting a role for brassinosteroid signaling in wheat inflorescence development.

During the inflorescence development time course in tetraploid Kronos presented here, 43.1% of genes were expressed in at least one timepoint, comparable to the 40.2% and 42.5% of genes expressed in similar inflorescence development time courses in the hexaploid wheat genotypes ‘Chinese Spring’ and ‘Kenong 9204’ when these reads were reanalyzed using the same mapping parameters and reference genome ^24,25^. Of these genes, 3,682 exhibited spike-dominant expression profiles (τ > 0.9). Among these genes were seven of ten SRS TFs, including the wheat ortholog of *six-rowed spike 2* (*HvVRS2*) that modulates hormone activity in the developing barley spike ^46^. Its expression profile in wheat, coupled with its association with spikelet number in an earlier study ^26^, suggests it plays a conserved role in wheat inflorescence development. It would also be interesting to characterize the function of four other SRS TFs that exhibit spike-specific expression profiles peaking towards terminal spikelet formation (Supplementary data 7). Ten of fifteen GRF TFs were expressed predominantly in spike tissues, including *TaGRF4* which improves regeneration efficiency in tissue culture when co-expressed with GIF cofactors^47^. The broadly similar, spike-specific expression profiles of genes in this family suggest other members may also contribute to meristem differentiation and inflorescence development (Supplementary data 7).

A subset of *WOX* TFs and *CLE* peptides exhibited dynamic and spike-dominant expression profiles across the time course, consistent with the differential regulation of *OsWUS, OsWOX3, OsWOX4*, and *OsWOX12* during panicle development in rice ^48^. The overexpression of *TaWOX5* (named *TaWOX9* in the current study) enhances wheat transformation and callus regeneration efficiency ^49^. Several other WOX TFs are co-clustered with this gene and exhibit similar expression profiles in the wheat inflorescence (Fig. 5), suggesting they may also be candidates to enhance regeneration efficiency (Fig. 5). Among CLE peptides, *TaCLV3* was negatively associated with spikelet number in a set of Chinese wheat landraces ^26^, consistent with its proposed role as a negative regulator of SAM size and activity in rice and maize ^50,51^.

Analyses of principal components and co-expression profiles indicate that the transition from the vegetative meristem to the double ridge stage is associated with major reprogramming of the wheat transcriptome (Fig. 1), consistent with an earlier study ^25^. Several TF families were enriched in module 1, characterized by high expression in the vegetative meristem before rapid downregulation after the double ridge stage, including eight of the nine RAV TFs in the consensus network. In Arabidopsis, the RAV genes *TEMPRANILLO1* (*TEM1*) and *TEM2* repress *FT* to prevent precocious flowering ^52,53^. In rice, the *TEM* orthologs *OsRAV8* and *OsRAV9* bind the promoters of *OsMADS14* and *Hd3a* to suppress the floral transition, indicating this function is conserved in monocots ^54^. The rapid downregulation of the wheat orthologs of these genes before double ridge formation, as well as homologs of *OsRAV11* and *OsRAV12* that act in reproductive patterning in rice ^54^, suggests this family may act as local repressors of meristem identity genes in the developing wheat spike.

There were also 26 TCP TFs clustered in module 1, including *TaTCP-A9* and *TaTCP-B9*, negative regulators of spikelet number and grain size in durum wheat ^55^. It is likely that other members of the TCP TF family also play roles as negative regulators of grain development. For example, *TaTCP-A17* and - *B17* are both downregulated during inflorescence development, are within 250 kbp of QTL for grain size, and are orthologous to genes associated with spikelet number variation in rice (Supplementary data 7). Eight TCP TFs clustered in different modules and were most highly expressed during spikelet meristem formation, including *TEOSINTE BRANCHED 1*, which integrates photoperiod signals to regulate spike architecture in a dosage-dependent manner ^56^, and a paralogous copy on chromosome 5B, *BRANCHED AND INDETERMINATE SPIKE*, that regulates spike architecture in barley ^57^. Four other uncharacterized TCP TFs with homology to *RETARDED PALEA1* exhibit spike-dominant expression profiles and would be promising candidates to characterize their role in inflorescence development in wheat (Supplementary data 7).

Although association and linkage mapping studies in wheat have described hundreds of QTL for agronomic traits, relatively few causative genes have been cloned and validated ^7^. Transcriptomic data can help prioritize candidate gene selection within a mapping interval based on spatial or temporal expression profiles ^58^. Furthermore, changes in transcription may indicate the presence of dominant or semi-dominant gain-of-function variants in *cis*-regulatory elements or of structural variation that confer changes in phenotype through modified expression profiles. Because of the functional redundancy of the polyploid wheat genome, such variants underlie the majority of cloned genes to date ^59^, including domestication alleles of *PPD1, VRN1* and *RHT1*, which clustered in different co-expression modules to their wild-type homoeologous allele (Fig. 4). Such divergent expression profiles, especially for those genes in close proximity to QTL for traits relating to grain number and grain size, might be strong candidates for allele mining to explore the extent of natural variation in wheat germplasm collections, and to engineer novel variation by targeted editing of *cis*-regulatory regions ^60^.

## Conclusions

Consensus and gene regulatory networks provide the means to analyze temporal transcriptomic datasets as a complementary approach to characterize functional pathways underlying wheat inflorescence development. The incorporation of higher resolution datasets at both the spatial and temporal levels within meristem tissues will build on these findings ^29^. Although reverse genetics will be required to validate the hypotheses generated from *in silico* network analyses, the integration of functional datasets from wheat and related species facilitates the identification of critical regulators ^61^. An improved understanding of the regulation of inflorescence development will help breeders combine superior alleles to drive increased grain number.

## MATERIALS AND METHODS

### Plant materials and growth conditions

All experiments were performed in the tetraploid *Triticum turgidum* L. subsp. *durum* (Desf.) var. Kronos (genomes AABB). Kronos has a spring growth habit conferred by a *VRN-A1* allele containing a deletion in intron 1 and carries the *Ppd-A1a* allele that confers reduced sensitivity to photoperiod ^62,63^. Plants were grown in controlled conditions in PGR15 growth chambers (Conviron, Manitoba, Canada) under a long day photoperiod (16 h light/8 h dark) at 23 □ day/17 □ night temperatures and a light intensity of ∼260 μM m^-2^ s^-1^. Developing apical meristems were harvested under a dissecting microscope using a sterile scalpel and placed immediately in liquid nitrogen. All samples were harvested within a one-hour period approximately 4 h after the lights were switched on (+/- 30 min) to account for possible differences in circadian regulation of gene expression. Approximately 20 apices were combined for each biological replicate of samples harvested at stages W1.0 (shoot apical meristem, SAM) and W2.0 (early double ridge, EDR) and approximately 12 apices for samples harvested at stages W3.0 (double ridge, DR), W3.25 (lemma primordia, LP) and W3.5 (terminal spikelet, TS) ^3^. Four biological replicates were harvested at each timepoint.

### RNA-seq library construction and sequencing

Tissues were ground into a fine powder in liquid nitrogen and total RNA was extracted using the Spectrum™ Plant Total RNA kit (Sigma-Aldrich, St. Louis, MO). Sequencing libraries were produced using the TruSeq RNA Sample Preparation kit v2 (Illumina, San Diego, CA), according to the manufacturer’s instructions. Library quality was determined using a high-sensitivity DNA chip run on a 2100 Bioanalyzer (Agilent Technologies, Santa Clara, CA). Libraries were barcoded to allow multiplexing and all samples were sequenced using the 100 bp single read module across two lanes of a HiSeq3000 sequencer at the UC Davis Genome Center.

### RNA-seq data processing

‘Kronos’ RNA-seq reads were trimmed and checked for quality Phred scores above 30 using Fastp v0.20.1 ^64^. Trimmed reads were aligned to the IWGSC RefSeq v1.0 genome assembly consisting of A and B chromosome pseudomolecules and unanchored (U) scaffolds not assigned to any chromosome (ABU) using STAR 2.7.5 aligner (outFilterMismatchNoverReadLmax = 0.04, alignIntronMax = 10,000) ^20,65^. Only uniquely mapped reads were retained for expression analysis. Transcript levels were quantified by featureCounts using 190,391 gene models from the ABU IWGSC RefSeq v1.1 annotations ^28,66^ and converted to Transcripts Per Million (TPM) values using a custom python script available from https://github.com/cvanges/spike_development/ (Supplementary data 1).

Raw RNA-seq reads for ‘Kenong9204’ and ‘Chinese Spring’ inflorescence development datasets were obtained from BioProjects PRJNA325489 and PRJNA383677 ^24,25^. RNA-seq reads were processed with Fastp as described above and aligned to the hexaploid ABDU RefSeq v1.0 genome assembly using the same methods and parameters. Transcript quantification and TPM were determined as above using the full ABDU IWGSC RefSeq v1.1 annotations.

RNA-seq reads and raw count data for each sample is available from NCBI Gene Expression Omnibus under the accession GSE193126 (https://www.ncbi.nlm.nih.gov/geo/).

### Transcription factors

There were 3,838 ABU gene models annotated as transcription factors that were grouped into 65 TF families per IWGSC v1.1 annotations ^28^. The following families were consolidated: “AP2” and “APETALA2”, “bHLH” and “HRT-like”, “MADS” and “MADS1”, “NFYB” and “NF-YB”, “NFYC” and “NF-YC”, and “SBP” and “SPL”, as well as “MADS2” and “MIKC”, which were consolidated into “MIKC-MADS”. After consolidation, there were 59 TF families. A previous study described he annotation of 201 MIKC-MADS box genes placed into 15 subclades ^17^. There were 30 MIKC transcription factors on the A and B genomes absent from the IWGSC TF list, which were added to this family. Investigations of the *CLE* and *WOX* gene families were based on the naming reported in Li et al., 2019b and Li et al. 2020b, with the addition of *TaWUSb* (*TraesCS2B02G775400LC*) to the *WOX* family, which was absent from these studies. In total, 3,861 TFs were included in this study (Supplementary data 2).

### Spike-dominant expression analysis

Expression data (TPMs) for two developmental studies were obtained from the Grassroots Data Repository (https://opendata.earlham.ac.uk/wheat/under_license/toronto/Ramirez-Gonzalez_etal_2018-06025-Transcriptome-Landscape/expvip/RefSeq_1.0/ByTranscript/) ^27,28^. The first dataset, in ‘Chinese Spring’, included samples from five tissue types at three timepoints (mean of two biological replicates) for 15 total tissue/stages ^27^. A second dataset from the variety ‘Azhurnaya’ comprised 209 unreplicated samples grouped into 22 “intermediate tissue” groups of various sizes^28^. Twelve samples overlapping with ‘Kronos’ spike samples were removed (tissue groups “coleoptile”, “stem axis”, and “shoot apical meristem”). For early spike tissue specificity analyses, the mean TPM expression of 15 ‘Chinese Spring’ tissues (n = 2) or the mean of 22 ‘Azhurnaya’ tissues (n ranging from 3 - 30) were compared to the ‘Kronos’ sampling stage with the highest mean expression (n = 4). Comparisons were made using the Tau (τ) tissue specificity metric where τ = 0 indicates ubiquitous expression and τ = 1 indicates tissue specific expression ^67,68^. A custom R script was used to calculate tissue specificity and is available at github.com/cvanges/spike_development. Genes which were expressed predominantly in ‘Kronos’ inflorescence tissues (τ > 0.9) were defined ‘spike-dominant’ whereas genes only expressed in ‘Kronos’ inflorescence tissues (τ = 1) were defined ‘spike-specific’ (Supplementary data 3).

### Principal Component Analysis (PCA), Differential Expression, and GO enrichment

PCA was performed in R using prcomp in the r/stats package v2.6.2 including all replications for each time point. PCA plots were generated with ggplot2 v3.3.2. Whole transcriptome PCA used read counts from all expressed gene models (n = 82,019) and TF PCA used expression of 2,874 expressed TFs. Randomized PCA distribution (Fig. S1) used independent random subsampling of 2,874 expressed genes without replacement. Principle component percent variation explained and eigenvalues from prcomp were used for comparisons between whole transcriptome PCA and TF-only PCA.

Pairwise differential expression was determined using both EdgeR v3.24.3 and DESeq2 v1.22.2 for robustness ^69,70^. Pairwise comparisons between consecutive timepoints were done using raw read counts for four biological replicates at each stage. Benjamin-Hochberg FDR adjusted *P*-values ≤ 0.01 was used as a stringent DE cut-off for both tools. Only genes DE using both tools were classified as pairwise DEGs (Supplementary data 5). Differential expression of ‘Chinese Spring’ and ‘Kenong9204’ inflorescence development datasets was also determined with raw read counts and EdgeR and DESeq2 using the same method as for the ‘Kronos’ dataset. Adjustments to DE tests were made to compare all four timepoints (6 pairwise comparisons) with two biological replicates in ‘Chinese Spring’ as well as the six timepoints (15 pairwise comparisons) with two biological replicates in the ‘Kenong9204’ datasets. For network analyses, a second DE test was included which reinforced longitudinal DE determination, an impulse model (ImpulseDE2, https://github.com/YosefLab/ImpulseDE2) was used for ‘Kronos’ data ^71,72^. Raw counts were used with default parameters and genes with Benjamin-Hochberg FDR adjusted *P*-values ≤ 0.05 considered differentially expressed. Functional annotation to generate GO terms for each high-confidence and low-confidence gene in the IWGSC RefSeq v1.1 genome was performed as described previously ^73^.

### Standard and consensus WGCNA network construction

Genes identified using pairwise differential expression (EdgeR and DESeq2) and ImpulseDE2 (22,566 genes total) were used for co-expression analyses. A standard co-expression network was built using the R package WGCNA v1.66 with the parameters: power = 20, networkType = signed, minimum module size = 30, and mergecutheight = 0.25 ^74^ (Supplementary data 7). Parallel coordinate plots were produced in R by normalizing raw read counts and visualized with ggparacoord (scale = ‘globalminmax’) in GGally (version1.5.0).

A consensus network was built using methods described in Shahan *et al*. (2018). In brief, 1,000 WGCNA runs were performed with 80% of genes randomly subsampled without replacement and random parameters for power (1, 2, 4, 8, 12, 16, 20), minModSize (40, 60, 90, 120, 150, 180, 210), and mergeCutHeight (0.15, 0.2, 0.25, 0.3). The final consensus network was built using an adjacency matrix – adj = number of times gene *i* is clustered with gene *j* / number of times gene *i* is subsampled with gene *j* – with parameters power = 6 and minModuleSize = 30 (Supplementary data 7). The consensus100 network was built by filtering the adjacency network for adj = 1 prior to network construction. Along with module assignments, we used the WGCNA package to find the connectivity of each gene with co-clustered genes (intramodularConnectivity.fromExpr()) and summarized module expression patterns (moduleEigengenes()). Python and R scripts for creating the adjacency matrix and consensus network are available at https://github.com/cvanges/spike_development. The Bioconductor package GeneOverlap was used to determine the overlap of module assignments between consensus and standard networks (http://shenlab-sinai.github.io/shenlab-sinai/) ^75^.

### Causal Structure Inference Network

Expression data (TPM) for 970 transcription factors retained in the consensus100 network was used to build a gene regulatory network using the Causal Structure Inference algorithm ^42^. Network construction used CSI in Cyverse with default parameters ^42^

### Conversion of wheat, rice, and barley gene IDs

Genes associated with wheat and rice spikelet number described in Wang et al., 2017b were identified from a previous set of annotated wheat gene models (ftp://ftp.ensemblgenomes.org/pub/plants/release-28/). To identify the corresponding IWGSC RefSeq v1.1 gene ID, each gene model coding sequence was extracted and used as a query in BLASTn searches against the IWGSCv1.1 ABU genome. Homologous gene pairs with > 99% identity to each query were considered spikelet number associated genes. Two previous studies reported genes DE during *H. vulgare* inflorescence development using IBSC_v2 annotations ^76,77^. Each barley gene model coding sequence was extracted and used as a query in BLASTn searches against the IWGSCv1.1 ABU wheat genome. Genes with percent identity > 90% were retained and considered orthologs of barley DEGs (HvDE).

### Enrichment analysis

Enrichment and depletion of genes among modules or DEG lists was determined using the cumulative distribution function of the hypergeometric distribution (systems.crump.ucla.edu/hypergeometric/).

### QTL proximity and definition of homoeologous pairs

Using a previously published meta-analysis of yield component QTL studies, we searched the IWGSCv1.1 genome for expressed genes in our timecourse within 500 kbp of 428 loci associated with yield component traits (kernel number per spike, thousand kernel weight, spikelet number) ^7^. Homoeologous gene pairs reported from Ramírez-González et al., (2018) ^28^ were used to determine co-expressed homoeologs.

## Acknowledgements

We thank Dr. Luis de Haro for assistance in RNA-seq library preparation and Dr. Cristobal Uauy for helpful input and discussion on project design and analysis. Work in this project was funded by the International Wheat Yield Partnership Grant IWYP76.

## Author contributions

SP and JD conceived of and designed the project. CV, JH and FT analyzed RNA-seq data. CV, SP and JD wrote the manuscript.

## Data Availability

All RNA-seq data have been deposited with the Gene Expression Omnibus (GEO) database under record number GSE193126. Processed expression and gene annotation information are provided as supplementary files.

## Additional Information

N/A

## Supplementary Files

Supplementary Data 1 – Expression levels (TPM) of 82,019 genes expressed in Kronos inflorescence transcriptome.

Supplementary Data 2 – Expression levels (TPM) of transcription factors during inflorescence development. TF families, subclade (for MIKC-MADS box TFs) and mean expression at each timepoint.

Supplementary Data 3 – Genes with spike-dominant expression profiles, including Tau specificity score, annotation, and TF family.

Supplementary Data 4 – Significantly enriched Gene Ontology terms associated with spike-dominant genes. Results for biological processes, cellular components and molecular functions are presented separately.

Supplementary Data 5 – Differential expression statistics for 82,019 genes expressed during inflorescence development. FDR-adjusted *P* and log_2_-fold change values are provided from ImpulseDE2 and all ten pairwise comparisons (DESeq2 + EdgeR).

Supplementary Data 6 – Significantly enriched Gene Ontology terms associated with genes differentially expressed in pairwise comparisons between consecutive timepoints. Results for biological processes, cellular components and molecular functions are presented separately.

Supplementary Data 7 – Information on 22,566 genes assembled into co-expression gene networks (standard WGCNA, consensus, consensus100) module assignments, intramodular connectivity (Kw), common gene name, and supporting evidence (TaSN, OsSN, TaDE, HvDE, QTL proximity, spike-dominant).

Supplementary Data 8 – Significantly enriched Gene Ontology terms associated with genes in each module of the consensus network. Results for biological processes, cellular components and molecular functions are presented separately.

Supplementary Data 9 – Significantly enriched Gene Ontology terms associated with genes in consensus100 module 9. Results for biological processes, cellular components and molecular functions are presented separately.

Supplementary Data 10 – Causal Structure Inference network (separate. xgmml file, Cytoscape compatible).

