## Supplemental Figures S1-S3 for "Transcriptional signatures of wheat inflorescence development"

^6^ Current address: Rothamsted Research, Harpenden, Hertfordshire, AL5 2JQ, UK.

*Corresponding author

Email addresses:

**Supplemental Figures 1 – 3**

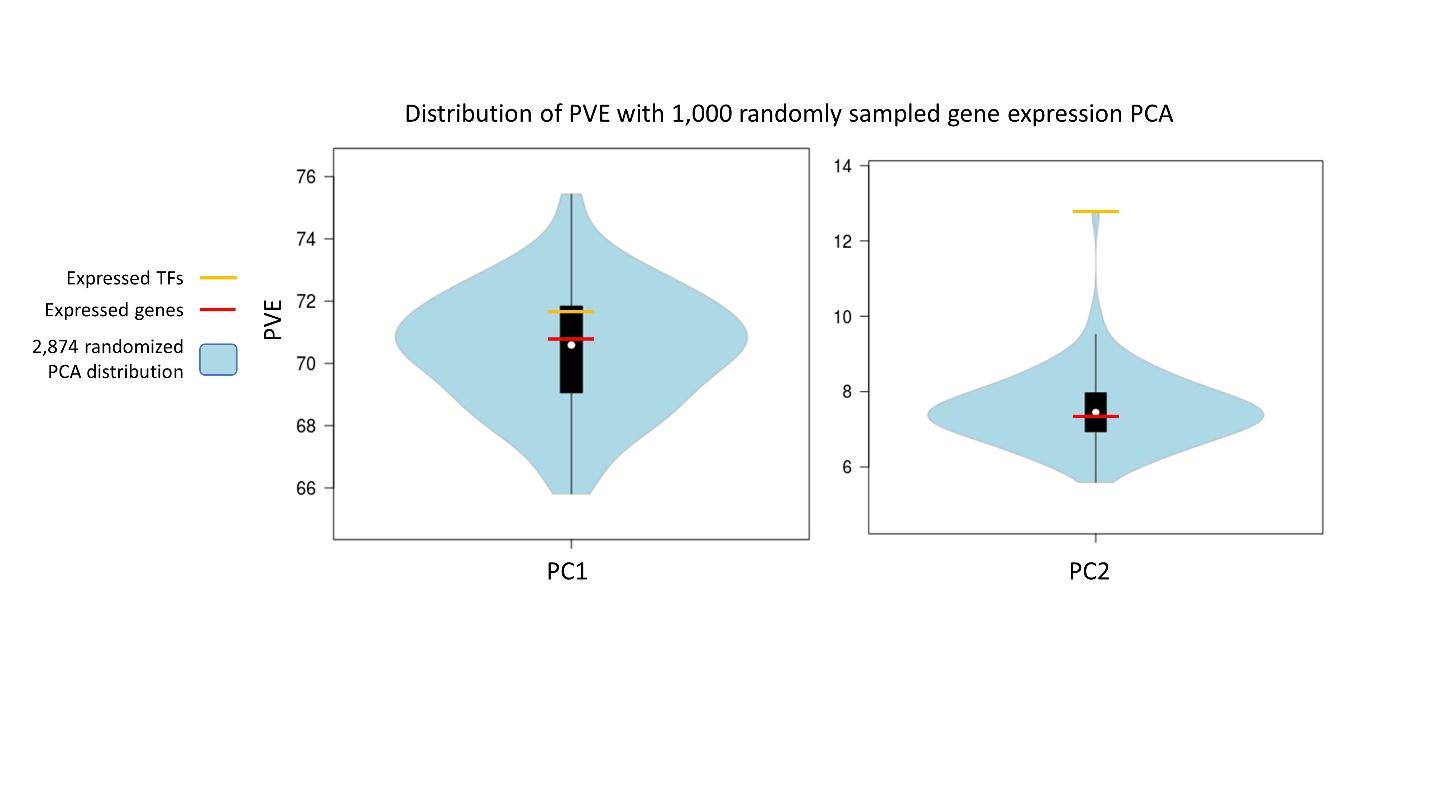

Figure S1: Percent variance explained by PC1 and PC2 considering expressed genes (n = 82,019, red), expressed transcription factors (n = 2,874, yellow), and sets of randomly subsampled genes (n = 2,874, blue). Violin plots represent distribution of PVE for PC1 and PC2 when 2,874 expressed genes are subsampled from all expressed genes 1,000 times. The variance explained by PC1 was only marginally greater compared to the whole transcriptome PCA, but PC2 captured a greater proportion of variance (12.77 PVE, Fig. S1) than when including the whole transcriptome (7.37 PVE). A one-sample t-test showed the PVE by PC2 of TFs was significantly higher than randomly subsampled genes of the same sample size from the whole dataset (P < 0.0001, df = 9,999, Fig. S1).

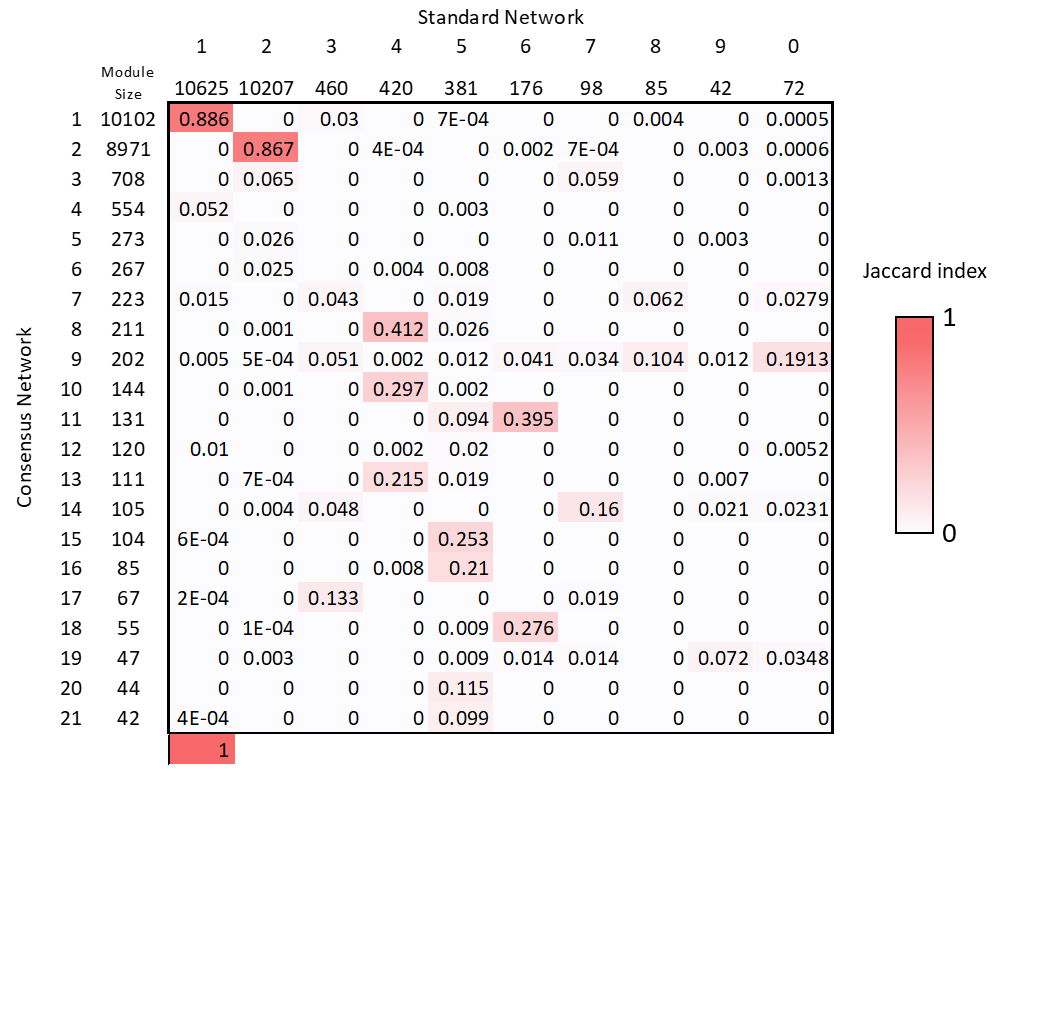

Figure S2: Comparison of common genes in consensus and standard co-expression network modules using the Jaccard index. Module sizes are listed for consensus (left) and standard network (above). Jaccard index scores range from 0 (No overlap) to 1 (identical gene lists).

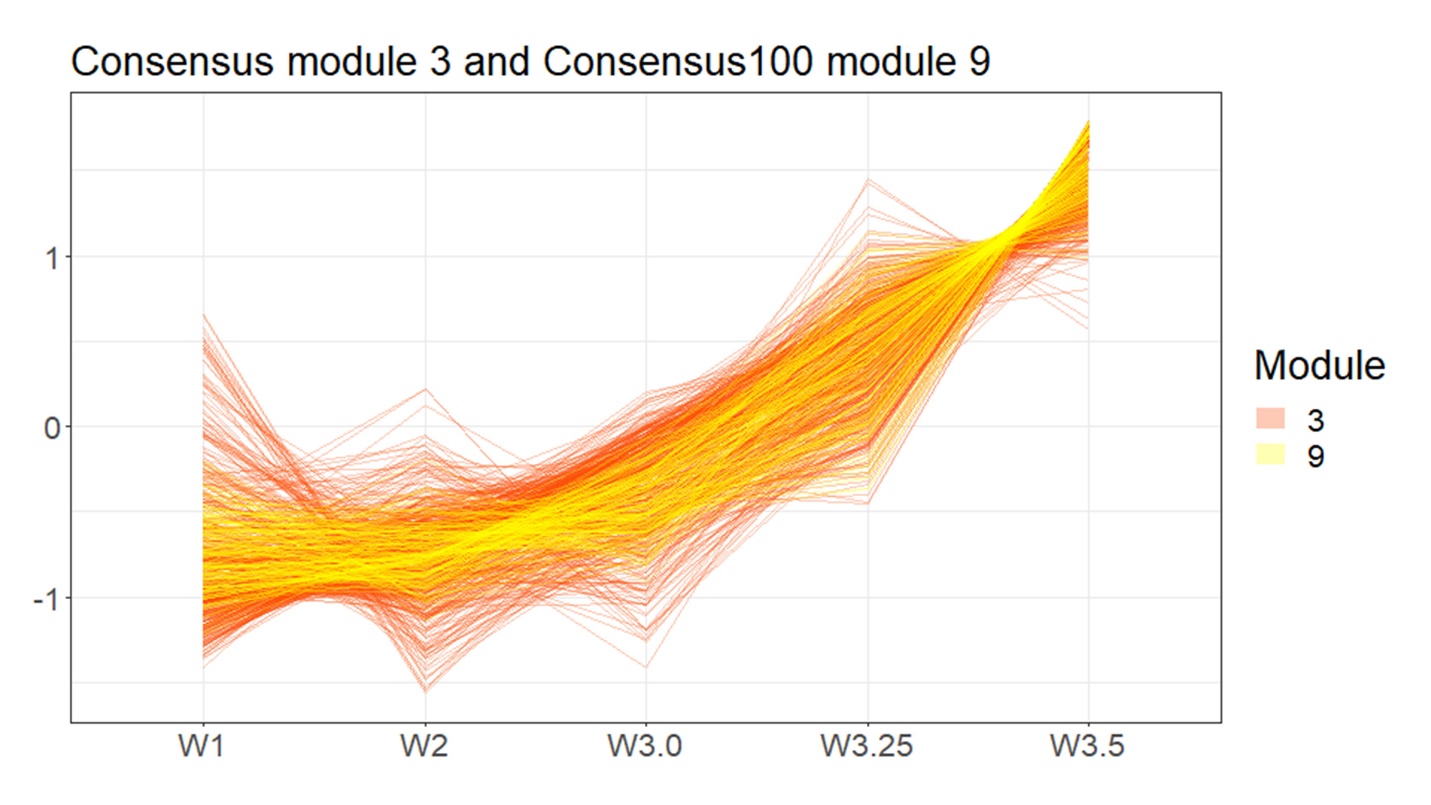

Figure S3: Expression profiles of all 708 genes in consensus network module 3 (orange) and the subset of 167 genes that were clustered in consensus100 module 9 (yellow).
